# Bayesian Inference of a Spectral Graph Model for Brain Oscillations

**DOI:** 10.1101/2023.03.01.530704

**Authors:** Huaqing Jin, Parul Verma, Fei Jiang, Srikantan Nagarajan, Ashish Raj

**Affiliations:** Department of Radiology and Biomedical Imaging, University of California San Francisco, USA San Francisco, CA; Department of Epidemiology and Biostatistics, University of California San Francisco, USA San Francisco, CA

**Keywords:** Bayesian, Connectomes, Magnetoencephalography, Spectral graph theory, Simulation-based inference

## Abstract

The relationship between brain functional connectivity and structural connectivity has caught extensive attention of the neuroscience community, commonly inferred using mathematical modeling. Among many modeling approaches, spectral graph model (SGM) is distinctive as it has a closed-form solution of the wide-band frequency spectra of brain oscillations, requiring only global biophysically interpretable parameters. While SGM is parsimonious in parameters, the determination of SGM parameters is non-trivial. Prior works on SGM determine the parameters through a computational intensive annealing algorithm, which only provides a point estimate with no confidence intervals for parameter estimates. To fill this gap, we incorporate the simulation-based inference (SBI) algorithm and develop a Bayesian procedure for inferring the posterior distribution of the SGM parameters. Furthermore, using SBI dramatically reduces the computational burden for inferring the SGM parameters. We evaluate the proposed SBI-SGM framework on the resting-state magnetoencephalography recordings from healthy subjects and show that the proposed procedure has similar performance to the annealing algorithm in recovering power spectra and the spatial distribution of the alpha frequency band. In addition, we also analyze the correlations among the parameters and their uncertainty with the posterior distribution which can not be done with annealing inference. These analyses provide a richer understanding of the interactions among biophysical parameters of the SGM. In general, the use of simulation-based Bayesian inference enables robust and efficient computations of generative model parameter uncertainties and may pave the way for the use of generative models in clinical translation applications.

## 1 Introduction

A key endeavor in the field of neuroscience is to uncover the relationship between the brain’s complex electrophysiological and functional activity, and its underlying structural wiring contained in white matter fiber projections [1, 2]. Functional activity between the grey matter regions is estimated with functional magnetic resonance imaging (fMRI), electroencephalography (EEG), and magnetoencephalography (MEG), while the structural wiring is assessed using diffusion tensor imaging (DTI) from MRI. The brain structure-function (SC-FC) relationship is then investigated using various data-driven and mathematical modeling-based techniques, assuming structural connectivity (SC) as a graph with different brain regions as graph nodes connected to each other via edges that are informed by the white matter fiber projections.

While both data-driven [3–17] as well as modeling techniques [18–28] have been employed to uncover the brain SC-FC relationships, mathematical models additionally provide insights into the underlying biophysics of brain activity. After fitting the model to empirical fMRI, EEG, and MEG data, the inferred model parameters can serve as biophysically interpretable markers of disease and brain states [29–34]. For example, Zimmermann et al. [33] demonstrated that the model parameters can predict cognition. However, the practical impact of model-based biomarkers of pathophysiology is hampered by two key challenges, described below.

### Lack of confidence bounds and posterior probabilities

An important goal of practical model fitting is to quantify how well a model’s parameters explain empirical neuroimaging data, and how confidently those estimates can be obtained. It is, therefore, necessary to capture their variability and find out all possible parameter settings compatible with the observed phenomena [35]. Bayesian inference is the established approach for achieving these goals, by making available the posterior distribution of parameters given the observations. Posterior distribution in turn provides rich information about how model parameters interact together, and quantifies the uncertainty of the model output - potentially critical for obtaining computational biomarkers in disease. Unfortunately, Bayesian inference methods have been proven to be quite challenging for most current computational models of brain activity.

### Tractability of model inference

We identify three issues limiting the tractability of Bayesian model evidence in the field. First, powerful sampling methods like Markov Chain Monte Carlo require extremely large samples, numbering in the hundreds of thousands. Most current models, like the coupled neural mass models (NMMs), are evaluated via time-consuming numerical integration techniques, which in turn impose a prohibitive computational burden on any sampling technique. Second, coupled NMMs involve large parameter spaces, i.e. number of internal parameters that must be jointly inferred, making full Bayesian inference impractical. Third, due to inherent non-linearity, the theoretical posterior density in even the simplest computational models is so convoluted, non-smooth, and non-convex that conventional optimization or MCMC sampling techniques encounter huge challenges. Many of these issues were highlighted in previous studies [36–38], and together they have ensured that hardly any Bayesian inference is performed in these settings.

In this paper, we present a novel way for Bayesian inference of computational models of neural activity, focusing specifically on the recently proposed spectral graph model (SGM), a linear biophysical generative model that can accurately capture the steady state wide-band power spectral density (PSD) as well as the spatial distribution of the alpha band power obtained from MEG [39]. We choose the SGM for the following reasons:

1. SGM involves a parsimonious set of global biophysically interpretable parameters; in our previous paper, we demonstrated that only 7 global, spatially-invariant parameters, each having distinct biophysical meaning, were sufficient to accurately capture empirical MEG PSD [40, 41]. This may be compared against previous models that have typically required substantially more spatially-varying parameters.
2. SGM explicitly estimates regional PSD and therefore can directly fit the frequency PSD obtained from MEG/EEG. Other models typically provide time-domain simulations only, and their spectral content is usually not a target of model fitting.
3. SGM is extremely fast to evaluate since its solution can be obtained in a closed-form in the frequency domain. Other models typically require lengthy time-domain simulations, which can be impractical in MCMC or other sampling techniques.

As a result of its linearity and closed-form evaluation without the need for long simulations, SGM parameter inference is far more tractable compared to non-linear neural mass models – where identifiability of model parameter is not guaranteed [36–38].

In our prior works we had estimated SGM parameters using a global optimization algorithm, the dual annealing method, as point estimates [39–41, 37], as a preferred alternative over continuous gradient descent-based minimization. However, since the objective function of estimating SGM parameter [39, 40] is non-convex, the annealing approach does not guarantee a global optimum, Moreover, a single point estimate from the annealing method is far from enough to uncover the underlying entire range and behavior of biophysical processes and to lead to new insights. On the contrary, Bayesian method allows the estimation of the full posterior distribution of the SGM parameters, which is necessary for biological interpretation. As a result, Bayesian method is more suitable for inferring the SGM parameters. However, the conventional Bayesian inference is challenging due to the fact that the theoretical posterior density of SGM parameters may be rather complicated which causes difficulty in sampling.

To circumvent the computational difficulty, we propose a novel method to perform Bayesian inference of the SGM parameters. The method approximates the posterior density of the SGM parameters by using a neural network model, which is trained through a simulation-based inference (SBI) framework [42]. Our main contribution is to show that this custom combination of SGM with SBI is exquisitely well-matched for estimating posterior distribution of generative model parameters.

This provides a far more appealing practical utility, which may be exploited in future clinical applications. Given its speed, this tool can be used to quickly infer posteriors of model parameters for a large number of subjects which can subsequently be used to identify parsimonious markers of disease and brain states. Finally, it allows us to benefit from the availability of an unbounded number of simulations, thereby helping overcome the critical issue of lack of large sets of empirical data in medical settings.

Using the SBI tool applied to the SGM model, we demonstrate that the model posteriors can accurately capture the empirical spatial distribution of alpha frequency band and PSD in MEG, and the inference of posteriors is substantially faster than the point estimate inference algorithm used in prior works. This combination of a fast and parsimonious forward model (SGM) with a fast neural network for posterior inference (SBI) is not currently available in the field of SC-FC mapping, and could constitute a critical advance in the applicability of computational models to practical scenarios.

## 2 Methods

### 2.1 Dataset

We study the resting-state Magnetoencephalography (MEG) data obtained from 36 healthy subjects (23 males, 13 females; 26 left-handed, 10 right-handed; mean age 21.75 years, age range 7-51 years) as also reported in Raj et al.’s study [39, 43]. Data collection procedure was describled in [39, 40]. All study procedures were approved by the institutional review board at the University of California at San Francisco and were in accordance with the ethics standards of the Helsinki Declaration of 1975 as revised in 2008. MEG recordings were collected for 5 minutes while the subjects were resting and had their eyes closed. Out of the 5-minute recording, a 1-minute snippet was chosen which was most noise free. MRI followed by tractography was used to generate the connectivity and distance matrices. The publicly available dataset consisted of processed connectivity and distance matrices, and PSD for every subject. MEG recordings were downsampled to 600 Hz, followed by a bandpass filtering of the signals between 2 to 45 Hz using `firls` in MATLAB [44] and generation of the static frequency PSD for every region of interest using the `pmtm` algorithm in MATLAB [44].

### 2.2 Spectral graph model

Spectral graph model (SGM) is a hierarchical, linear, analytic model of brain oscillations, which has a closed-form solution in the Fourier frequency domain via the eigen-decomposition of a graph Laplacian [39–41]. A typical SGM has two model layers, a mesoscopic layer for the local neuronal subpopulations of every brain region and a macroscopic layer for the long-range excitatory neuronal subpopulations of the whole brain. SGM is briefly described below, and detailed illustrations can be found in the supplementary document and in prior publications [39, 40].

SGM is characterized by eight parameters, which include the excitatory and inhibitory time constants *τ_e_*, *τ_i_* and neural gains *g_ee_*, *g_ei_* and *g_ii_* at the mesoscopic level, and long-range excitatory time constant *τ_G_*, coupling constant *α*, speed *v* at the macroscopic level. The neural gain *g_ee_* is kept as 1 to ensure parameter identifiability [39], so the parameters of interest to be inferred in SGM are **s** = (*τ_e_*, *τ_i_*, *α*, *v*, *g_ei_*, *g_ii_*, *τ_G_*)^*T*^. Given the signals with *N* regions of interest (ROIs) in the time domain is [*x*_1_(*t*),…, *x_N_*(*t*)]^*T*^, the closed-form solution of SGM is obtained in the Fourier domain:

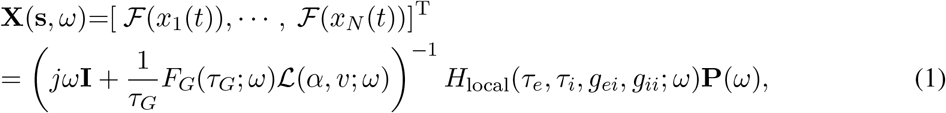

where *ω* is the angular frequency, **X**(**s**, *ω*) is a vector of the Fourier transformation, or equivalently the PSD, of the macroscopic signal over all brain regions of interest at frequency *ω*, 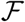 is the Fourier transformation, 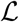 is the complex Laplacian, *H*_local_ is the mesoscopic model’s transfer function, **P**(*ω*) is the input noise spectrum, and *F_G_*(*ω*) is the Fourier transform of a Gamma-shaped neural response function, given as 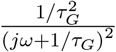. This response function is governed by the characteristic long-range excitatory time constant *τ_G_*, and the function is intended to serve as a lumped model of various processes, including dendritic arborization, somatic conductance, synaptic capacitance, etc [39].

### 2.3 Simulation-based inference for SGM

Simulation-based inference (SBI) is a powerful tool for the inference of large complex statistical models that have been extensively applied in many areas of science and engineering [45–47]. We adapt the SBI method for SGM parameter estimation and inference (referred to as SBI-SGM). Let **X**(**s**, Ω) = {**X**(**s**, *ω*)}_*ω*∈Ω_ be the model output PSD in dB scale[48] where Ω is the set of the frequency points we used and it contains 40 equally spaced frequencies in the range 2-45 Hz in the manuscript. *G*{**X**(**s**, Ω)} is a monotonic transformation that standardizes the PSD across the frequency into a z-score; standardizes the regional distribution of alpha band power (i.e. summation of PSD from 8-12 Hz); and finally concatenates both into a single vector. Here and throughout the text, we present the PSD in dB scale. In our SBI-SGM framework, we assume the data model is

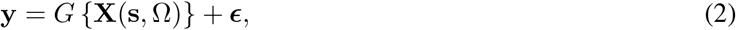

where ***ϵ*** ~ *N*(**0**, *σ*^2^**I**) is additive i.i.d. noise with standard deviation (SD) *σ*.

The random noise in (2) captures the biological noise, artifacts, and measurement error in the MEG data. Without this random noise, the target posterior density is discontinuous, which is difficult to estimate due to the well-known Gibbs phenomenon [49]. Adding this random noise to the model results in a smooth posterior distribution of SGM parameters, which can be accurately approximated by a neural network [50].

Similarly, we define the observed samples **y**_*o*_ = *G* {**X**_MEG_(Ω)} where **X**_MEG_(Ω) is the observed MEG PSD.

Since the parameters in the SGM model are assumed to be bounded to satisfy biological constraints, bounded priors are typically adopted for them, which causes difficulties for posterior sampling with SBI [51]. To address this issue, we re-parameterize the parameters so that the posterior sampling can be performed on the real line. More specifically, let 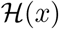 be a scaled logit transformation function [52] defined as

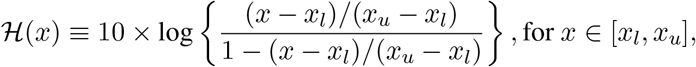

where *x_l_* and *x_u_* are lower and upper bounds of variable *x*, respectively. Slightly abusing the notation, let 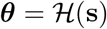, where 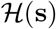 represents the values of function 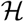 applied on each element of **s**.

Under a Bayesian framework, we are interested in the posterior distribution of ***θ*** given **y**, particularly the Bayesian credible interval of ***θ***, which captures the uncertainty of the SGM parameters. To obtain the credible interval, we estimate the posterior distribution of ***θ*** through the SBI procedure [42]. The density of **y** is denoted by *p*(**y**|***θ***) following (2), which is a multivariate Gaussian density function. We impose a multivariate Gaussian prior *π*(***θ***) on ***θ***. The posterior density is *q*_**Φ**_(***θ***|**y**) ∝ *π*(***θ***)*p*(**y**|***θ***), where **Φ** is the unknown parameters that determine the posterior distribution. Instead of obtaining the posterior density for the SGM parameters **s** directly, we first derive the posterior density for ***θ***, which results in the target posterior distribution through a Jacobian transformation [53].

We use a deep learning architecture, namely neural spline flow (NSF) [47], to model the functional form of *q*_**Φ**_, where **Φ** is the parameters in the deep learning network. The dimension of **Φ** increases with the number of network layers in NSF, and when the dimension of **Φ** approaches infinity, *q*_**Φ**_ approaches the true posterior distribution. When the deep learning architecture is given, **Φ** is the only unknown parameter in *q*_**Φ**_(***θ***|**y**). Hence estimating the posterior density is equivalent to estimating **Φ**. Now note that the true posterior distribution maximizes **E**[log{*q*_**Φ**_(***θ***|**y**)}], where the expectation is taken with respect to **y** and ***θ***, we propose to obtain an estimator for **Φ** through maximizing the empirical version of **E**[log{*q*_**Φ**_(***θ***|**y**)}], that is 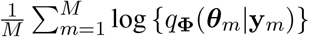, where the samples **y**_*m*_ and ***θ***_*m*_, *m* = 1,…, *M* are the simulated realizations of **y** and ***θ*** based on *p*(**y** |***θ***) and *π*(***θ***), respectively.

To obtain the posterior density for **s** given the observed sampled from the empirical PSD of MEG data ***y***_*o*_ = *G* {**X**_MEG_ (Ω)}, we can feed ***y***_*o*_ in the neural network and obtain the estimated posterior distribution *q*_**Φ**_(***θ***|**y**_*o*_) with the estimated parameter 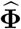. The target posterior distribution of **s** is 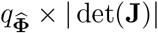 where **J** is the Jacobian matrix, i.e., **J** = *∂****θ***/*∂***s**. We illustrate the details of obtaining the posterior distribution of ***θ*** in Algorithm 1, which contains a simulation step and an optimization step.

#### Algorithm 1 Posterior estimation with re-parameterization

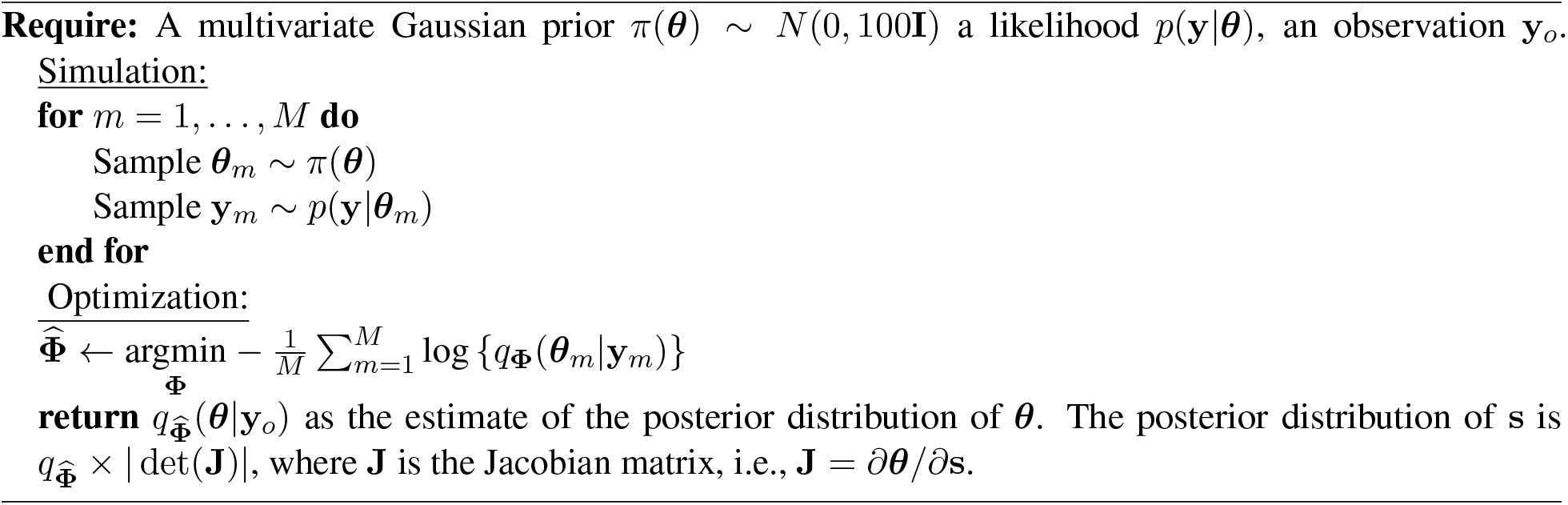

### 2.4 Implementation details

For each subject, we use their MEG data from 68 cortical regions according to the Desikan–Killiany atlas [54] to obtain the posterior samples of the SGM parameters **s**. Under this atlas, we obtain a 68-region **X**(**s**, *ω*) at frequency *ω*, and the dimension of *G* {**X**(**s**, Ω)} is 2788.

We implement SBI-SGM using the sbi package in Python (https://www.mackelab.org/sbi/) [55], where the hyperparameters in the original SBI algorithms are adopted as the default values provided in the package. We discuss the choice of the standard deviation of the noise *σ* and the number of simulation samples in <u>Simulation</u> step in Algorithm 1 in the next section.

In SBI-SGM, we adopt an average template structural connectome created via openly available diffusion MRI data obtained from the MGH-USC human connectome project (HCP) for training a universal posterior mapping from observation to the posterior distribution using Algorithm 1. After obtaining a trained posterior density for each observed **y**_*o*_, we draw a posterior sample of SGM parameters, denoted by 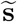. We then obtain 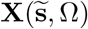 using (1). Finally, we construct the standardized PSD and spatial distribution of the alpha band PSD as a *G* transformation of 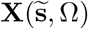, where the function *G* is defined in (2). We perform this posterior sampling process 1000 times to obtain a set of posterior samples of the standardized PSD and spatial distribution of the alpha band PSD for each observed **y**_*o*_. The pipeline of SBI inference for SGM is presented in Figure 1.

**Figure 1:**
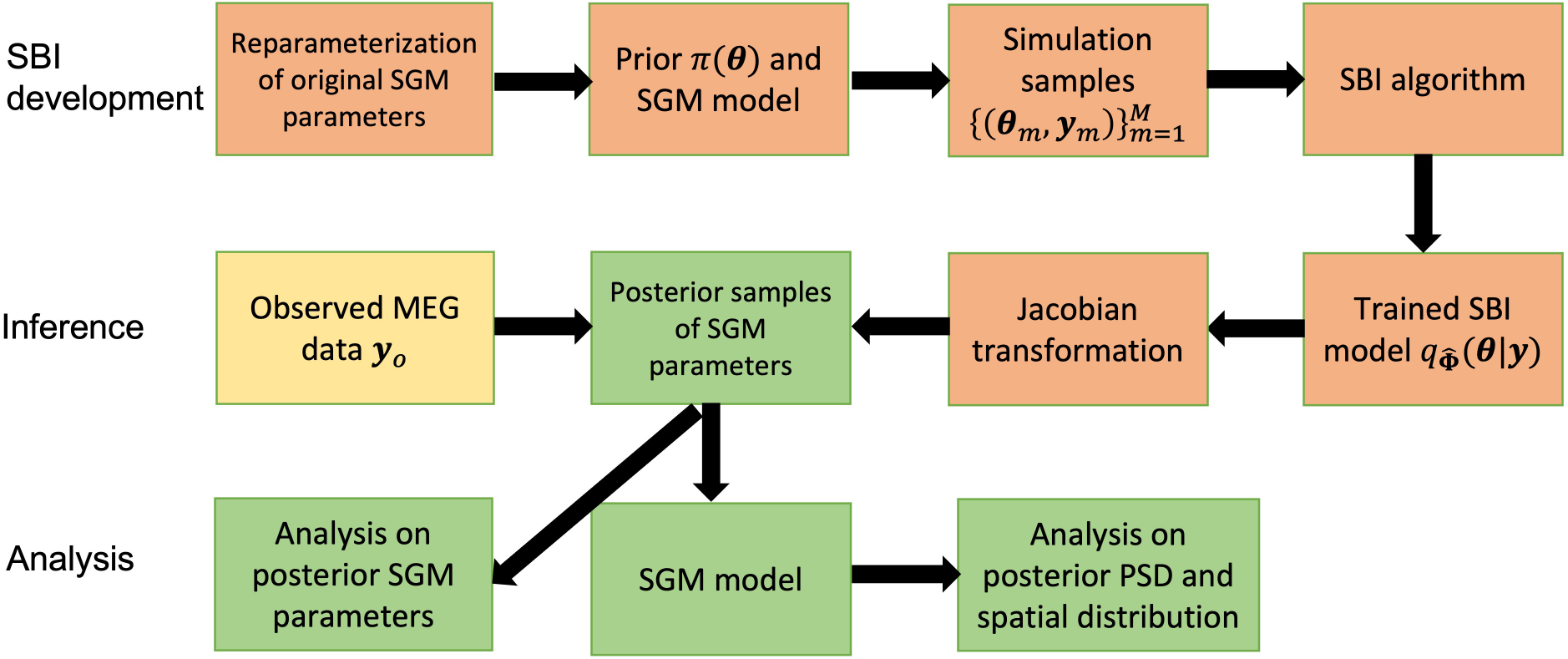
The pipeline of SBI inference for SGM.

We compare the performance of SBI-SGM with the performance of the annealing SGM approach [39–41], namely Ann-SGM, on our MEG data. The details of the annealing implementation can be found in [40]. SGM model assumes the parameters have finite supports as the ones listed in Table 1. In Ann-SGM, three different bounds are evaluated for *g_ei_* and *g_ii_* sequentially, and the largest bounds that satisfy the stability condition defined by [41] are chosen in the subsequent estimation. In SBI-SGM, the largest bounds in Table 1 are adopted for parameters (*g_ei_*, *g_ii_*) and we only retain the posterior samples of the SGM parameters within the stability boundary defined in [41]. For the other five parameters, SBI-SGM uses the same bounds as Ann-SGM does.

**Table 1:** SGM parameters and bounds for the parameter estimation for SBI and annealing.

| Name | Symbol | Lower/upper bound |
| --- | --- | --- |
| Excitatory time constant | $\tau_e$ | [0.005 s, 0.03 s] |
| Inhibitory time constant | $\tau_i$ | [0.005 s, 0.2 s] |
| Long-range connectivity coupling constant | $\alpha$ | [0.1, 1] |
| Transmission speed | $v$ | [5 m/s, 20 m/s] |
| Alternating population gain | $g_{ei}$ | [0.001,0.7], [0.001,0.5], [0.001,0.4] |
| Inhibitory gain | $g_{ii}$ | [0.001,2.0], [0.001,1.5], [0.001,1.5] |
| Graph time constant | $\tau_G$ | [0.005 s, 0.03 s] |
| Excitatory gain | $g_{ee}$ | n/a |

## 3 Results

### 3.1 Adding random noise to the SGM improves the reconstructing accuracy of the PSD

For each subject, we obtain the reconstructed PSD by taking the mean of the posterior samples of the PSD. We then study how the change in noise variation affects the performance of SBI-SGM in reconstructing the observed PSD. We compare the median Pearson’s correlation between the reconstructed PSD and the observed PSD from MEG. Specifically, for each ROI, we calculate the correlation between the reconstructed PSD and observed PSD from MEG. We then average the correlations over all ROIs and obtain the median of this average correlation over 36 subjects. In this study, the number of simulations samples is fixed at 100, 000 in the <u>Simulation</u> step in Algorithm 1 and the standard deviation of *ϵ* varies from 0 to 3.2. We report the mean results over 10 repetitions. Note that when *σ* = 0, there is no random noise added.

Figure 2A shows that compared to the model without random error (when the *σ* = 0), adding random noise in (2) significantly reduces the reconstruction errors. This result is consistent with our theoretical conclusion that adding random noise results in a smooth posterior density which can be accurately approximated by a neural network. The Pearson’s correlation between the reconstructed and the observed PSD increases when *σ* < 1.6 and starts to decrease after *σ* reaches 1.6, when the signal-to-noise ratio is not sufficiently large for the SBI-SGM to recover the observed PSD. In practice, we suggest choosing *σ* in [0.8, 2.0], which yields satisfactory performance with over 0.9 correlation between reconstructed and observed PSD. For all the following experiments, we fix *σ* = 1.6.

**Figure 2:**
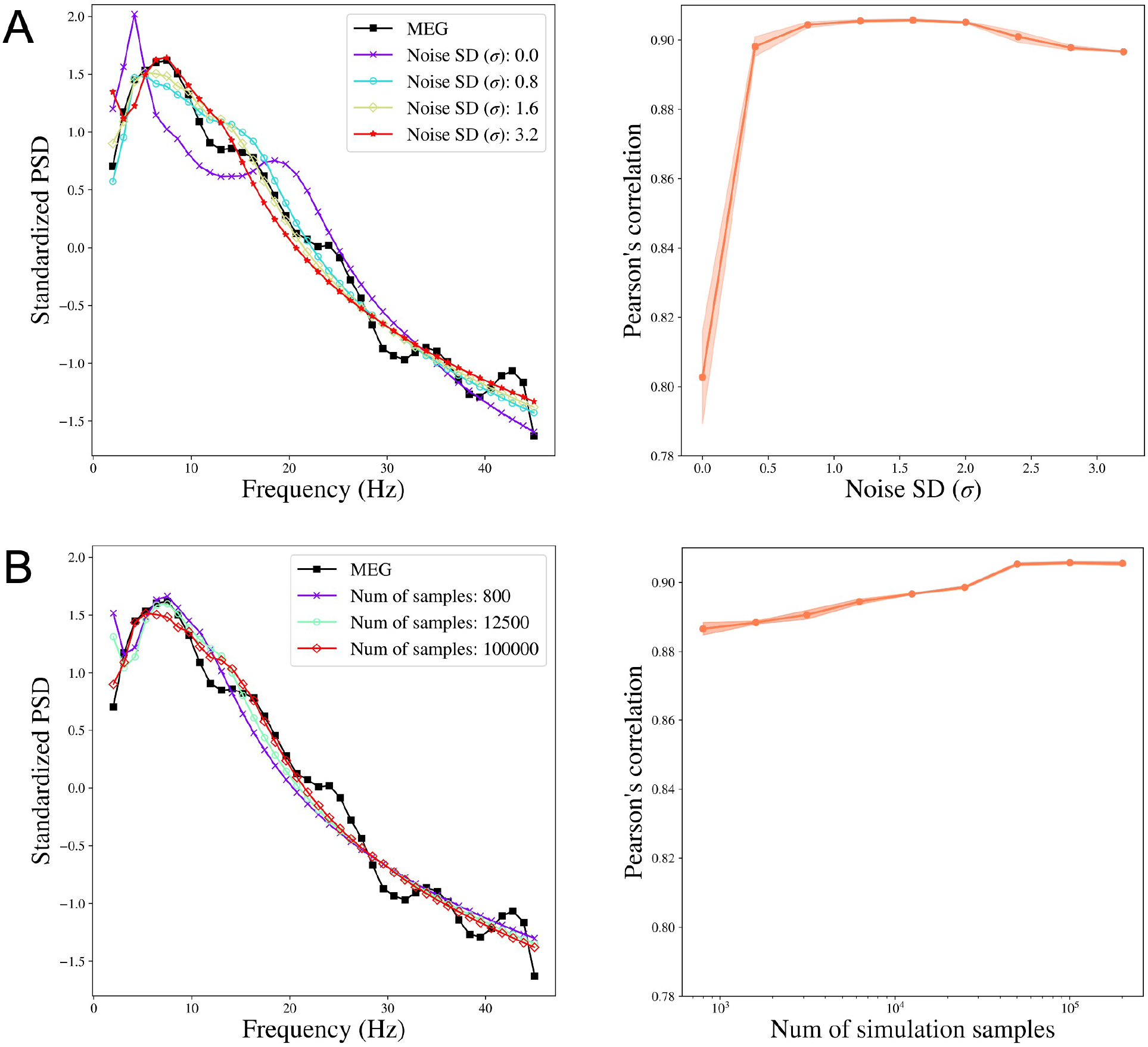
The performance of SBI-SGM when varying the noise SD and the number of simulation samples. **A:** Left: Median standardized power spectral density (PSD) obtained from MEG and SBI-SGM with different noise SDs. Right: Change of Pearson’s correlation between reconstructed average PSD and the observed PSD when varying noise SDs. The red shadow indicates its 95% confidence interval. **B:** Left: Median standardized PSD obtained from MEG and SBI-SGM with different number of simulation samples. Right: Change of Pearson’s correlation between reconstructed average PSD and the observed PSD when varying the number of simulation samples. The red shadow indicates its 95% confidence interval.

### 3.2 Increasing the number of simulation samples improves the SBI-SGM fit

We also investigate how the performance of SBI-SGM changes with the number of simulations in <u>Simulation</u> step in Algorithm 1. Figure 2B shows that a larger number of simulations yields a better SBI-SGM fit with a higher correlation between reconstructed and observed PSD. As indicated by the right panel of Figure 2B, the changes in Pearson’s correlation are not very notable and it varies from 0.887 to 0.906. However, when considering the standardized PSD curves for different numbers of samples, a clear trend is observed that typically larger sample size leads to a better fit visually. It is also worth noting that the performance of SBI-SGM is stable after the number of simulations reaches 100, 000. Therefore, we choose 100, 000 simulations in the <u>Simulation</u> step in Algorithm 1 in the subsequent analyses.

### 3.3 Results from two representative MEG data

We show the results from two representative subjects whose Pearson’s correlations between reconstructed PSD and the observed PSD are the top two closest to the median correlation across 36 subjects in one experiment. To make it representative, we repeat the SBI-SGM procedure 10 times and choose the experiment which yields an overall correlation closest to the mean level in the 10 repetitions for the analysis.

The posterior samples of the seven parameters as well as the PSD for two subjects are displayed in Figures 3A and B. In the left panels, we compare the posterior density of SBI-SGM with the point estimate from Ann-SGM. We can observe multiple modes from the posterior densities. For *τ_e_*, *v*, and *g_ii_*, the point estimates from Ann-SGM are close to one of the modes of the posterior distributions, while the estimates of the rest of the parameters from Ann-SGM are far away from the posterior modes from SBI-SGM.

**Figure 3:**
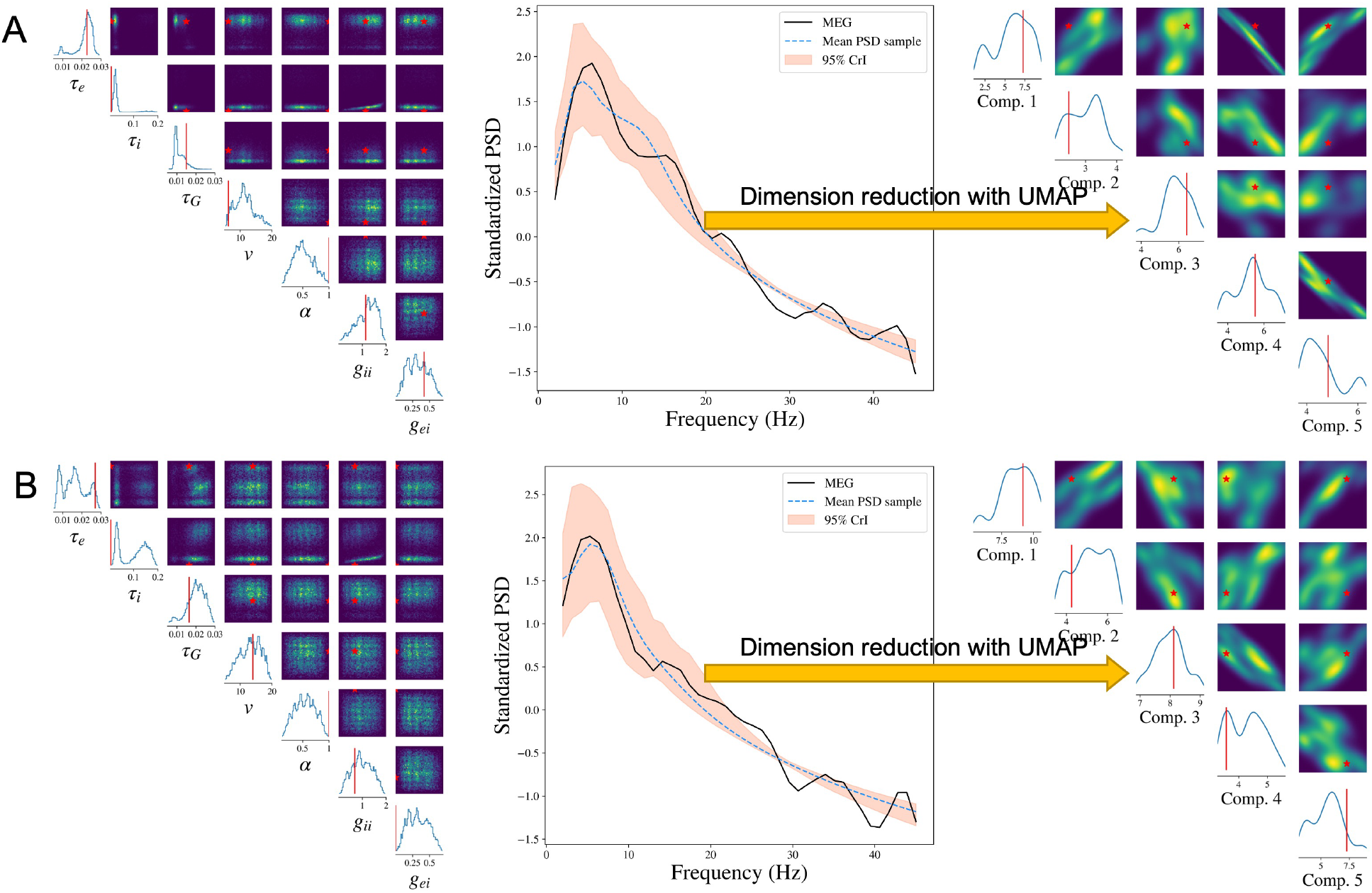
Result analysis from SBI-SGM for two representative MEG data. **A/B:** Left: Posterior density of 7 parameters for one subject in the MEG dataset. The red vertical line and red star indicate the location of the point estimate from the annealing algorithm. Middle: Posterior mean PSD and the 95% credible interval for the subject. The black curve indicates the observed average PSD. Right: Density estimations and observed values of low-dimensional representations after mapping raw PSDs to a 5-dimensional embedding manifold with the uniform manifold approximation and projection (UMAP). The red vertical line and red star indicate the location of the representation for observed MEG data in the manifold.

The middle panels of Figure 3 shows the posterior mean and 95% credible interval (CrI) of the PSD from SBI-SGM. In each subject, the 95% CrI covers the observed PSD at low frequencies (lower than 20 Hz), which is consistent with the fact that SGM can successfully recover the low-frequency PSD [40].

Moreover, we project the reconstructed and observed PSDs and map them onto a 5-dimensional manifold using the uniform manifold approximation and projection (UMAP) method proposed by [56]. As shown in the right panels of Figure 3, the projection of the observed PSD falls within the support of the projection of the reconstructed PSDs in the manifold, which further validates our Bayesian inference [57].

### 3.4 Cohort level analysis of MEG datasets

In SBI, the variability of the posterior distribution exists due to the randomness of the simulated samples in the <u>Simulation</u> step and the randomness in the posterior sampling procedure using the trained posterior distribution. We evaluate the robustness of SBI-SGM in 10 repetitions. In Figure 4A, we show the median of the reconstructed PSDs over 36 subjects for each repetition, the PSD Pearson’s correlation is changed between [0.905, 0.907] (shown in the caption). The results indicate that SBI-SGM is robust throughout the repetitions. We choose an experiment that yields a correlation closest to the mean level in the 10 repetitions in the subsequent analyses.

**Figure 4:**
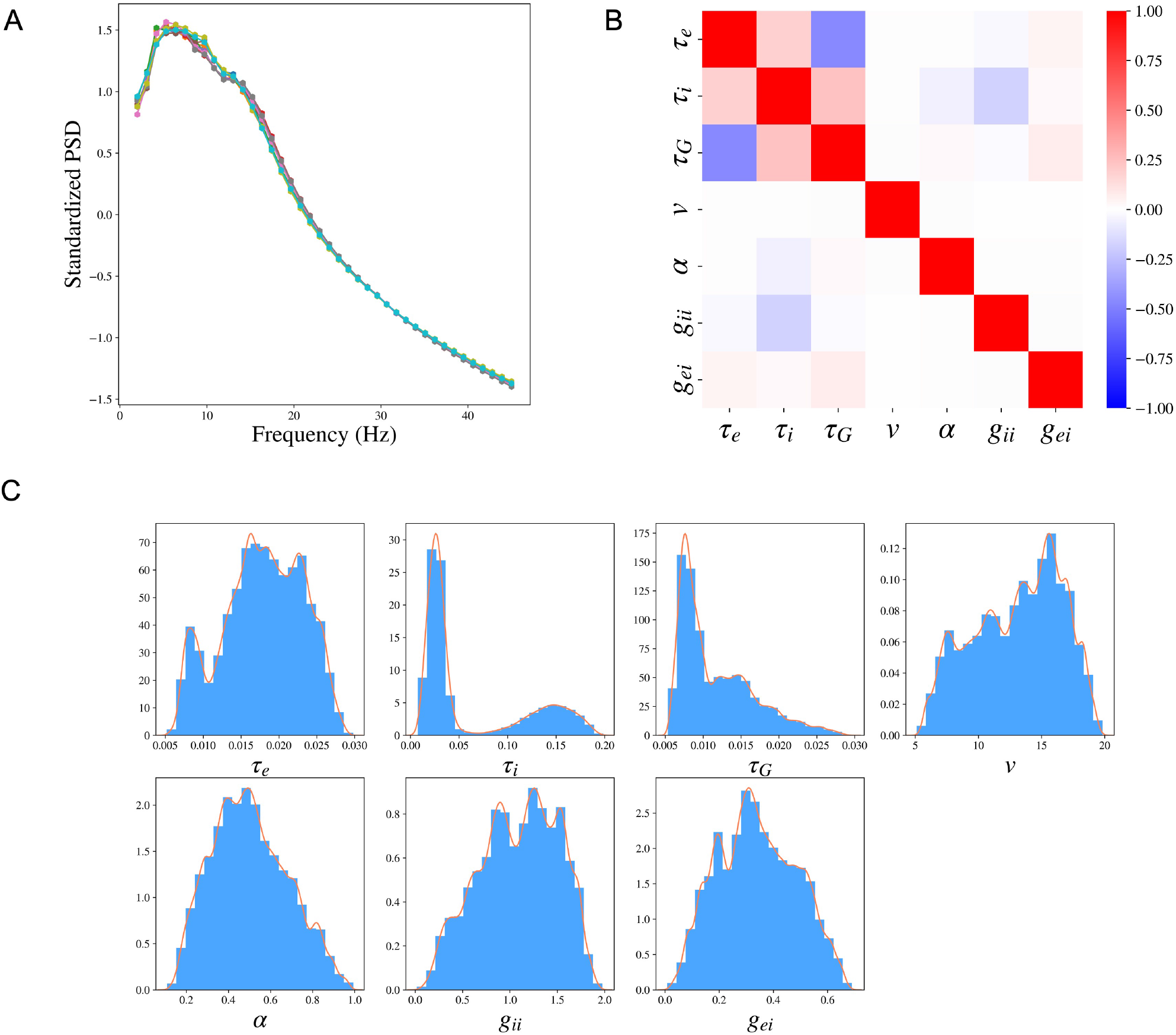
Cohort level SBI-SGM across the 36 MEG datasets. **A:** The median standardized power spectral densities (PSDs) obtained from SBI-SGM in 10 repetitions under noise SD 1.6 and number of simulation samples 100, 000. The correlations between reconstructed and observed PSDs are between 0.905 and 0.907 in the 10 repetitions. **B:** Partial correlation between each pair of parameters averaged over 36 subjects. **C:** Histograms and the corresponding kernel density estimations of the posterior SGM parameters.

We analyze the posterior samples of the SGM parameters from SBI-SGM across 36 MEG data. We first study the pair-wise correlation between the SGM parameters using the partial correlation method [58], which examines the correlation between any given two parameters after removing the effect from other parameters, Figure 4B shows the pair-wise partial correlation averaged over 36 subjects. As shown in Figure 4B, speed *v* has no correlation with the other parameters. The two time constants *τ_e_*, *τ_i_* have weak positive correlation, and the graph time constant *τ_G_* shows moderate negative correlation with the exhibitory time constant *τ_e_* and small positive correlation with the inhibitory time constant *τ_i_*. Figure 4C shows the distribution of the pooled posterior samples of the SGM parameters over 36 MEG data. Among the seven SGM parameters, the posterior distributions of *τ_i_*, *τ_G_* are highly concentrated, which indicates their variabilities across different subjects are small. The histogram of the inhibitory time constant *τ_i_* presents a second peak around 0.15s. The speed *v* has the highest density round 15 m/s.

We further investigate whether the SGM model in (1) can generate the observed PSDs. We generate 1000 SGM parameters from the prior distribution of **s**, and obtain simulated PSDs through (1). We then compare the simulated PSDs with the observed ones. To facilitate the visualization, we utilize the UMAP method to project the simulated samples of PSDs and observed PSDs to a 2-dimensional embedding manifold. As shown in Figure 5, in the embedded manifold, all the observed projections fall within the projections of the simulated samples, which indicates that the SGM model captures the generating mechanism of the observed PSD, and therefore is a reasonable likelihood of the data.

**Figure 5:**
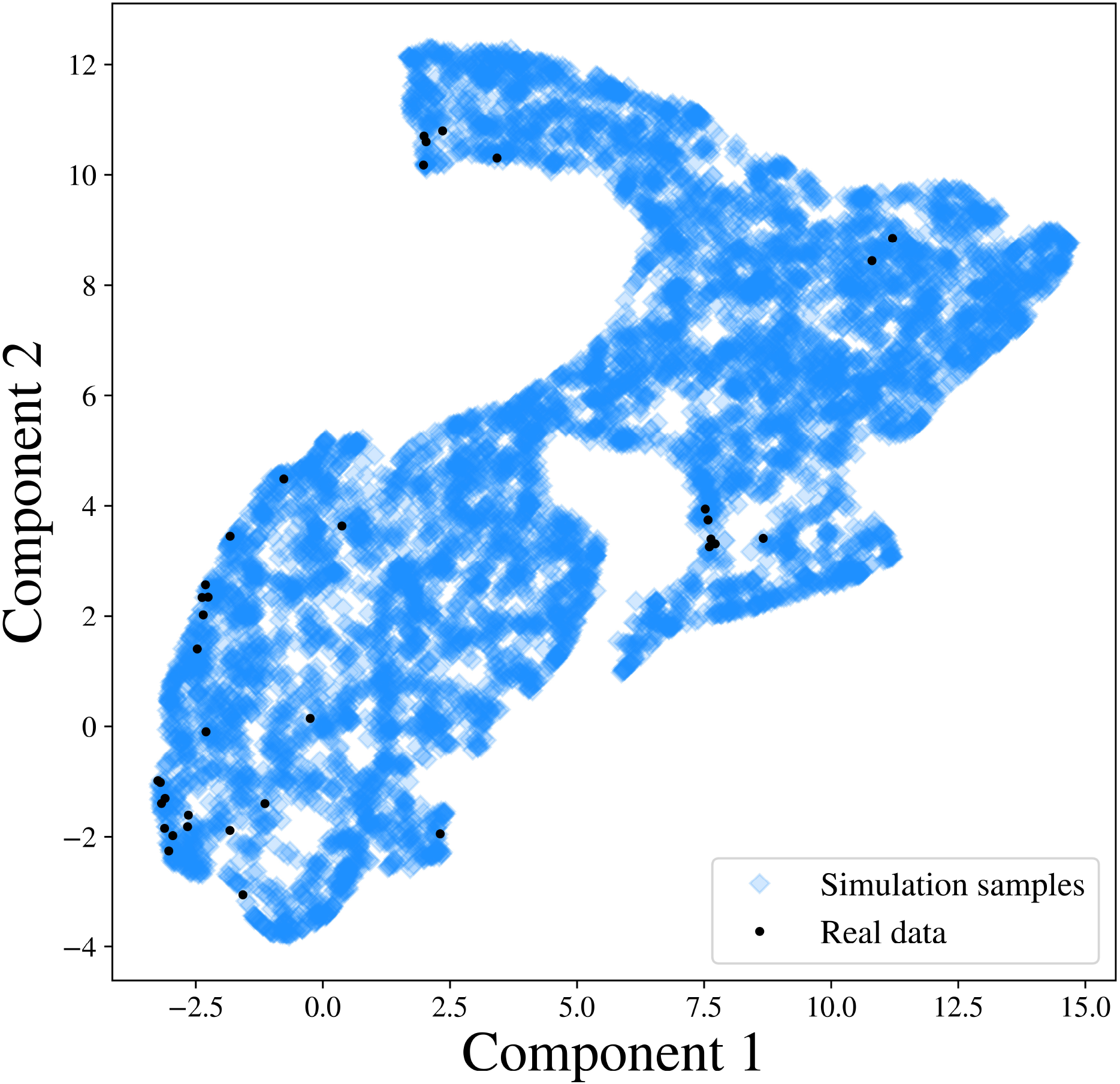
PSD representations after mapping the simulated PSDs and 36 observed PSDs to a 2-dimensional embedding manifold with the UMAP.

### 3.5 SBI-SGM and Ann-SGM comparision

We compare SBI-SGM with Ann-SGM. In Figure 6A, we show the correlations between the reconstructed and observed PSDs and the correlations between reconstructed and observed spatial distributions of the alpha band PSD resulting from SBI-SGM and Ann-SGM. We also perform statistical tests on the difference between the results from the two inference methods. Specifically, we calculate the Pearson’s correlations for each ROI between the reconstructed and observed PSDs and take the average across ROIs. Furthermore, we obtain the spatial correlation as the inner product between the reconstructed and observed spatial distribution of the alpha band PSD weighted by **D** + *w***I** where **D** is the row degree normalized structural connectivity matrix, **I** is the identity matrix, *w* is an empirical weight, and we adopt *w* = 10 as suggested by [37].

**Figure 6:**
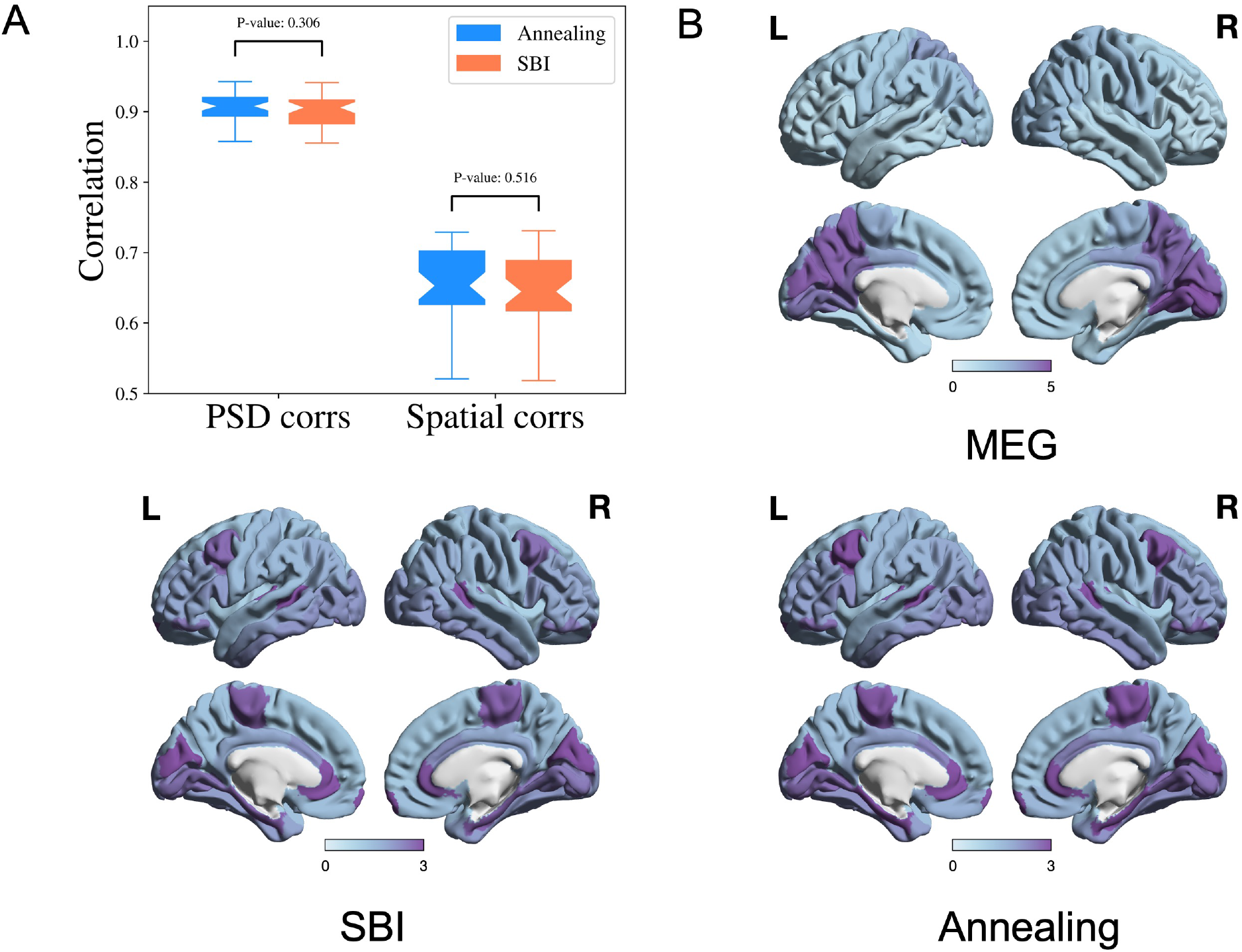
Performance of SBI and annealing on SGM is comparable. **A:** Pearson’s correlation of PSD from each ROI and spatial correlation for the alpha frequency band. P-values are from two-sample t-tests. **B:** Comparison of the observed and reconstructed spatial distributions from the SBI and annealing algorithms of the alpha frequency band, averaged over all the subjects

As shown in Figure 6A, SBI-SGM gives similar average correlation and spatial correlation as Ann-SGM does with insignificant p-values from two-sample t-tests. In Figure 6B, we observe very similar spatial distributions of the alpha band PSD from SBI-SGM and Ann-SGM, and both of them are similar to the observed one from MEG data.

One notable advantage of SBI-SGM is that it is much faster than Ann-SGM. When a template connectome is given, SBI-SGM provides a universal posterior distribution that can be applied to all the 36 MEG data after one training. SBI-SGM takes approximately 2 hours to accomplish the Bayesian inference on SGM parameters for all subjects. On the other hand, Ann-SGM takes about 8 hours for each subject and the algorithm needs to repeat for each subject. Parallel computing can further improve the computational speed of SBI-SGM. In conclusion, SBI-SGM has a similar performance as Ann-SGM on recovering observed PSD and spatial distribution from the alpha band but is much more computationally efficient.

## 4 Discussion

Models with complex and stochastic simulators have been extensively applied in many areas of science and engineering [59]. In neuroscience, such computational models are typically built via incorporating biological mechanisms and hypothetical intuitions to explain the observed phenomena inferred from the neuroimaging data [35]. These models involve several free parameters that are required to be compatible with the observed phenomena. Due to the complexity of neural models and neural data, the determination of the free parameters generally relies on computation-intensive optimization routines like grid search [60], genetic algorithm [61] or simulated annealing [39, 40].

However, these algorithms are far from meeting the needs of the neuroscience community, as they can only provide a single point estimate of the free parameters, and make it difficult to incorporate prior knowledge about related neural processes. In neural models, it is always desirable to find out not only the best, but all parameter settings compatible with the observed data. The variability of the parameters under the observation can provide more insights into the neural models and processes [35, 62]. Moreover, neural model parameters, e.g. of SGM, typically have biological meaning, hence their inference must accommodate the underlying biological mechanisms and their constraints, in order to avoid unreasonable solutions. Using the prior knowledge of these biological quantities can not only increase the optimization efficiency but also robustify the inferred models. Most importantly, the practical applicability of model fitting demands a solid assessment of the confidence bounds and variability associated with fitted parameters - something quite lacking in current methods. Due to these reasons, full Bayesian inference of posteriors is preferable to point estimates. However, the intricacy of the neural models typically results in intractable or complicated likelihood which makes the likelihood-based inference inaccessible.

Luckily, the SBI approach fills this gap by bypassing the evaluation of the likelihood function and giving the posterior samples directly. The results presented in this study have highlighted the key ways in which the proposed combination of SGM and SBI is exquisitely well-suited to the task of model inference of neural systems. First, the parsimony of SGM obviates a key weakness of SBI, which typically prefers to infer a small set of parameters [42, 47]. For this reason, SBI may be challenging for coupled non-linear models such as NMMs and the Virtual Brain [63] which consist of a potentially large set of parameters. Second, SBI requires a large number of forward-model evaluations to generate enough simulation samples for training, which would render large coupled NMMs unfeasible [24, 64, 27, 26], but this is far less problematic for SGM due to its fast forward evaluation. Third, SBI requires far fewer empirical samples compared to simulation samples [47, 42], which is an important consideration in real data-poor medical settings. Lastly, while the training of the neural network requires a high upfront cost involving numerous simulations, the trained SBI model can be applied almost instantaneously to new empirical data directly, which enhances the practical utility and amortizes the computational cost of fitting by front-loading the simulation effort.

We were able to show that the SBI-SGM framework gives speedy estimates of the full posterior distribution, achievable in a matter of seconds per subject. Using these posteriors, point estimates, e.g. mean or mode of the posterior, can be quickly produced, which we showed has comparable performance to prior point estimation methods like dual annealing, at a fraction of the computation time. Lastly, our posterior analysis showed that the model parameters were generally weakly correlated, implying that all of them are required to obtain model outputs that match the spectral and spatial patterns obtained from empirical MEG. This is a crucial finding since it suggests that we can identify unique markers of diseases and brain states in the form of inferred SGM parameters.

### 4.1 Relationship to previous works

For models closely related to SGM, such as the non-linear neural mass models or the dynamic causal models (DCM), Bayesian inferencing algorithms such as variational Bayes have been used previously. DCM employs variation Bayes to obtain effective functional connectivity [65, 66]. A key difference is that DCM is primarily used to obtain effective connectivity from smaller networks and that these connectivities are obtained from second-order statistics such as cross-spectra using spectral DCM [67]. In contrast, SGM directly computes the PSD rather than individual elements of the second-order effective connectivity matrix. SGM instead employs an explicit structure-based model, where the inter-regional connectivity comes directly from the measured structural connectome. In this manner, SGM is better suited for SBI than DCM, since the latter would be required to infer an entire matrix of effective connectivities, in addition to other regional or global parameters.

The key challenges with inferring parameters of coupled non-linear neural mass models are that they require time-consuming simulations. These models exhibit bifurcations yielding discontinuities in the model solutions [37], and parameter identifiability is not guaranteed [36]. These challenges have been discussed in detail elsewhere [38, 37]. SGM overcomes these challenges by providing a closed-form solution that can be simulated within seconds, and by consisting of only a parsimonious set of global model parameters.

Another potential way to conduct Bayesian inference for SGM is Markov chain Monte Carlo (MCMC) methods [68] as SGM has a closed-form solution in the Fourier frequency domain. However, even under an explicit frequency domain solution, the likelihood function of SGM can be complicated [41], which hampers analyzing the properties of the posterior density. Moreover, MCMC methods require a long burn-in step to reach the equilibrium distribution and samples from the equilibrium distribution are correlated. These properties make sampling from MCMC rather time-consuming for SGM. In addition, the computational cost of MCMC methods can not be amortized which means the time-consuming MCMC procedure needs to be run anew for each observation, regardless of prior observations. A previous MCMC-based inference was unable to capture the spectral features using a nonlinear neural mass model [38]. In comparison, SBI is more flexible than MCMC methods. Due to the powerful neural network, it can easily handle the complicated likelihood function. More importantly, SBI is trained with simulation samples that help to reduce the requirements of real data. Once the model is trained, it can be applied to new observations without retraining. Therefore, compared with MCMC methods, SBI may be preferable for practical Bayesian inference.

It is worth noting that while this paper focuses on SGM, the SBI approach can be a robust and efficient alternative for parameter estimation of any complex generative model, e.g. above-mentioned coupled neural mass or DCM models. The key trade-off involves whether upfront simulation of a large number of forward model runs is practical and whether there is a compelling use case for achieving rapid inference of an unseen observation.

### 4.2 Limitations of the current approach

In the current inference procedure, our simulator outputs include the regional PSD and the spatial distribution of alpha band power - together they form a relatively high-dimensional feature space. While Algorithm 1 is capable of handling this, the high dimensionality of output features increases the computational burden and causes difficulty in learning useful information with neural networks from the data. Although we reported some basic diagnostics in Figure 4 to verify the validity of our inference, the high-dimensional output hampers more extensive posterior diagnostics. Possible workarounds to deal with this issue include extracting some key features from the PSD and spatial distribution manually or embedding a neural network to learn the key features automatically. More experiments are required in this direction. Another limitation is the large number of simulation samples required in SBI-SGM, which slows the inference procedure and increases the computational burden. The number of required simulation samples can be dramatically reduced with multi-round inference [46] via focusing the training on a particular observation. Although the trained model loses the generality for other observations, it can be very useful when we are only interested in one specific observed dataset.

### 4.3 Potential applications and future work

In clinical practice, it can sometimes be even more important to know how accurate our estimate is than simply to know the best point estimate [69]. For example, using only point estimates, it can be difficult to compare computational biomarkers from different cohorts. Even if two cohorts have very different values of the biophysical parameters, no statistically robust conclusion can be drawn without knowing the uncertainty of those estimates. In such cases, SBI-SGM will be extremely helpful as it gives the posterior distributions of the parameters which fully captures the uncertainty of estimates. With posterior distributions, credible intervals and other measures of uncertainty can be easily obtained. This can also be used to obtain population-level parameters that are homogeneous across a population despite the individual variability, which can aid in establishing the descriptive validity of models like SGM [70]. Lastly, it can also be used to obtain time-varying posteriors of model parameters that can capture the fast temporal fluctuations in MEG, as has been done previously using point estimates [41].

## Data availability

The code and processed datasets for this work can be found in this github repository: https://github.com/JINhuaqing/SBI-SGM.

## Acknowledgments

This work was supported by NIH grants R01NS092802, R01NS183412, R01AG062196, R01AG072753, R01EB022717, R01DC013979, R01NS100440, R01DC176960, R01DC017091, K25AG071840, UCOP-MRP-17-454755, Alzheimer’s Association grant AARFD-22-923931, and an industry research contract from Ricoh MEG Inc. The template Human Connectome Project (HCP) connectome used in the preparation of this work was obtained from the MGH-USC HCP database (https://ida.loni.usc.edu/login.jsp). The HCP project’s MGH-USC Consortium (Principal Investigators: Bruce R. Rosen, Arthur W. Toga and Van Wedeen; U01MH093765) is supported by the NIH Blueprint Initiative for Neuroscience Research Grant; the National Institutes of Health grant P41EB015896; and the Instrumentation Grants S10RR023043, 1S10RR023401, 1S10RR019307. Collectively, the HCP is the result of efforts of co-investigators from the University of Southern California, Martinos Center at Massachusetts General Hospital (MGH), Washington University, and the University of Minnesota.

## Supplementary

### Spectral graph model

#### Notation

All the vectors and matrices are written in boldface and the scalars are written in normal font. The frequency *f* of a signal is specified in Hertz (Hz), and the corresponding angular frequency *ω* = 2*πf* is used to obtain the Fourier transforms. The connectivity matrix is defined as **C** = *c_jk_*, where *c_jk_* is the connectivity strength between regions *j* and *k*, normalized by the row degree.

### Mesoscopic model

Given region *k* out of *N* regions, we denote the local excitatory signal as *x_e_*(*t*), local inhibitory signal as *x_i_*(*t*), and the long-range macroscopic signals as *x_k_*(*t*). Combining the decay of individual signals, coupling of excitatory and inhibitory signals as well as input white Gaussian noise, the evolution models of *x_e_*(*t*) and *x_i_*(*t*) are:

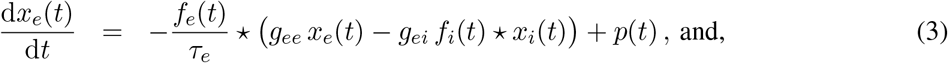

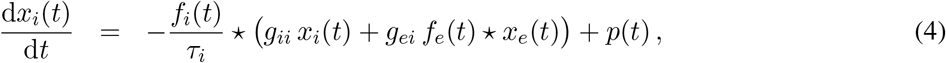

where *f_e_*(*t*) and *f_i_*(*t*) are the ensemble average neural impulse response function, ⋆ stands for convolution, *p*(*t*) is input noise, parameters *g_ee_*, *g_ii_*, *g_ei_* are neural gain terms, and parameters *τ_e_*, *τ_i_* are characteristic time constants, which are shared for every region *k*. We assume Gamma-shaped *f_e_*(*t*) and *f_i_*(*t*) as

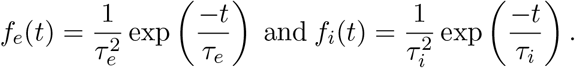

### Macroscopic model

Accounting for long-range connections between brain regions, the macroscopic signal *x_k_* is assumed to conform to the following evolution model:

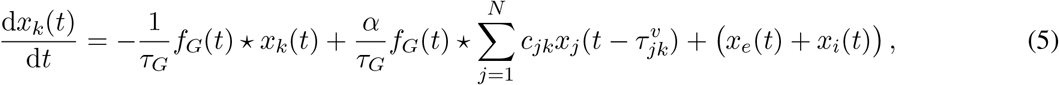

where, *τ_G_* is the graph characteristic time constant, *α* is the global coupling constant, *c_jk_* are elements of the connectivity matrix, 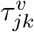 is the delay in signals reaching from the *j^th^* to the *k^th^* region, *v* is the cortico-cortical fiber conduction speed with which the signals are transmitted. The delay 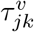 is calculated as *d_jk_*/*v*, where *d_jk_* is the distance between regions *j* and *k* and *x_e_*(*t*) + *x_i_*(*t*) is the input signal determined from Equations (3) and (4).

SGM only includes 8 global parameters as listed in Table 1. The neural gain *g_ee_* is kept as 1 to ensure parameter identifiability. Thus, there are only 7 parameters required to be estimated to determine SGM.

### Closed-form model solution in the Fourier domain

A salient feature of SGM is that it provides a closed-form solution of brain oscillations under the frequency domain. Let 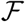 be the Fourier transform at angular frequency *ω* = 2*πf*. Note that the mesoscopic models for different regions share the same parameters, therefore, without loss of generality, we can drop the subscript *k*.

The solutions for *x_e_*(*t*) and *x_i_*(*t*) under the frequency domain are

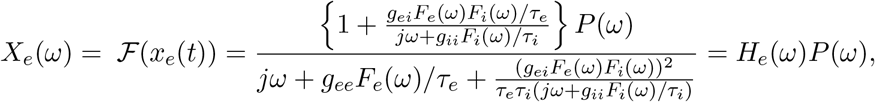

and

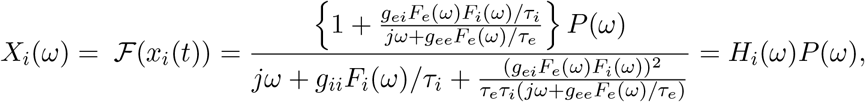

where *P*(*ω*), *F_e_*(*ω*), *F_i_*(*ω*) are the Fourier transform of *p*(*t*), *f_e_*(*t*) and *f_i_*(*t*) at angular frequency *ω*.

We define the complex Laplacian matrix 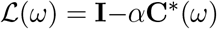 where 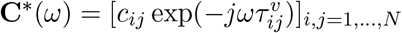.

The solution of the macroscopic signals at a angular frequency *ω* is

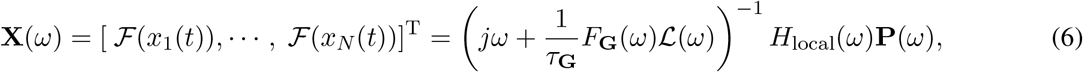

where *H*_local_(*ω*) = *H_e_*(*ω*) + *H_i_*(*ω*).

As SGM provides a closed-form solution **X**(*ω*), we can compare the modeled and empirical power spectra to estimate the global parameters.

